# GAISHI: A Python Package for Detecting Ghost Introgression with Machine Learning

**DOI:** 10.64898/2026.01.31.703038

**Authors:** Xin Huang, Josef Hackl, Harvinder Pawar, Martin Kuhlwilm

## Abstract

**Summary:** Ghost introgression is a challenging problem in population genetics. Recent studies have explored supervised learning models, namely logistic regression and UNet++, to detect genomic footprints of ghost introgression. However, their applicability is limited because existing implementations are tailored to tasks in their respective publications, but not available as user-friendly software implementations. Here, we present GAISHI, a Python package for identifying ghost introgressed segments and alleles using multiple machine learning algorithms and demonstrate its usage in different introgression scenarios.

**Availability and implementation:** GAISHI is available on GitHub under the GNU General Public License v3.0. The source code can be found at https://github.com/xin-huang/gaishi.

## Introduction

Introgression is a process leading to the presence of genetic material in a lineage that originates from another lineage, often termed ghost introgression where the donor lineage is unknown (Huang et al. 2025a). Genomic footprints consistent with ghost introgression have been reported in various species such as bonobos, gorillas, and hickories (Kuhlwilm et al. 2019; Pawar et al. 2023; Zhang et al. 2024). These investigations have been enabled by several methods based on summary statistics and probabilistic models that infer introgression signatures without requiring genomes from the donor population (Plagnol and Wall 2006; Vernot and Akey 2014; Vernot et al. 2016; Browning et al. 2018; Skov et al. 2018; Setter et al. 2020; Huang et al. 2022; Macià and Skov 2025), often centered on gene flow from archaic hominins into modern humans.

Recent advances in machine learning have further increased interest in applying these approaches, particularly deep learning, to population genetic inference (Schrider and Kern 2018; Huang et al. 2024). Several supervised machine learning-based implementations, including ArchIE with logistic regression and IntroUNET with UNet++, have been explored for detecting segments or alleles resulting from ghost introgression (Durvasula and Sankararaman 2019; Ray et al. 2024). However, these implementations are tailored to specific demographic models, limiting their applicability to other scenarios, and can be sensitive to implementation details, which may affect performance (Huang 2024; Hackl and Huang 2025). To address these issues, we developed GAISHI (version 1.0.0), a Python package for machine learning-based genomic analysis to identify introgressed sites and haplotypes.

### Implementation

GAISHI is implemented in Python and built on two machine learning libraries—scikit-learn and PyTorch (Paszke et al. 2019; Pedregosa et al. 2011). It generates training data via msprime simulations under demographic models specified in the Demes format (Baumdicker et al. 2022; Gower et al. 2022). Currently, GAISHI supports three supervised learning models: logistic regression, extra-trees classifiers (Geurts et al. 2006), and UNet++ (Zhou et al. 2019). Extra-trees is a tree-ensemble method, whereas UNet++ is a convolutional neural network architecture. GAISHI uses logistic regression and extra-trees classifiers to predict introgressed segments, and UNet++ to predict introgressed alleles.

For logistic regression and extra-trees classifiers, GAISHI converts either phased or unphased genotype data in VCF format (Danecek et al. 2011) into feature vectors computed from user-selected statistics. Supported statistics include pairwise Euclidean distances between genomes from the reference and target populations, pairwise Euclidean distances within the target population, the number of private mutations in the target population, per-sample allele frequency spectra in the target population, and the *S*\* statistic computed from the target genotype matrix. For the pairwise Euclidean distance features, users can additionally include the computation of distributional summaries of all pairwise distances, including the minimum, maximum, mean, median, variance, skewness, and kurtosis. Here, the reference population denotes the population that did not receive introgressed material, whereas the target population denotes the population that received introgressed material (Huang et al. 2025a).

For UNet++, GAISHI directly utilizes genotype matrices for training and inference. To accommodate variable sample sizes and enforce a fixed input shape for UNet++, GAISHI upsamples genotype matrices along the sample axis. When a desired number of samples is specified, additional columns are generated by sampling existing samples with replacement until the specified number is reached. The upsampled matrix is constructed by appending these duplicated samples to the original matrix. Genotype matrices are then sorted along the sample axis to impose a consistent ordering, because convolutional neural networks are sensitive to the spatial arrangement of the input (Huang et al. 2024). For the target population, all pairwise Euclidean distances between samples are computed, and a seriation algorithm is applied to obtain a permutation that places genetically similar samples next to each other. The target genotype matrix is reordered by this permutation, producing a structured, similarity-preserving layout. For the reference population, a cross-population distance matrix is computed between each target sample and each reference sample. A linear sum assignment problem is solved on this distance matrix to obtain a one-to-one matching that minimizes the total distance across matched pairs. Reference samples are then reordered according to the matched order, aligning the reference genotype matrix to the already-sorted target matrix.

For labeling, GAISHI extracts per-sample introgressed genomic intervals recorded in the msprime tree sequences. To label a simulated sequence as introgressed or non-introgressed, GAISHI uses two user-defined thresholds: an introgressed-proportion threshold (default = 0.7) and a non-introgressed-proportion threshold (default = 0.3). Let *p* denote the proportion of the simulated sequence covered by introgressed genomic intervals. If *p* is greater than or equal to the introgressed-proportion threshold, the sequence is labeled as an introgressed segment. If *p* is smaller than or equal to the non-introgressed-proportion threshold, the sequence is labeled as a non-introgressed segment. Simulated sequences with *p* between the two thresholds are discarded. In addition, an allele is labeled as introgressed if its genomic position falls within an introgressed interval extracted from the tree sequence; otherwise, it is labeled as non-introgressed.

### Examples

To demonstrate the usage of GAISHI, we applied it to three demographic models with different introgression scenarios (Figure 1): ArchIE_3D19, BonoboGhost_4K19, and HumanNeanderthal_4G21 (Durvasula and Sankararaman 2019; Kuhlwilm et al. 2019; Gower et al. 2021). ArchIE_3D19 models introgression from a source population that diverged 12,000 generations ago into the target population 2,000 generations ago with an admixture proportion of 0.02. BonoboGhost_4K19 models ghost-to-bonobo introgression 20,000 generations ago from a ghost population that diverged 140,000 generations ago, also with an admixture proportion of 0.02. HumanNeanderthal_4G21 models Neanderthal introgression into modern Europeans (CEU) 1,896 generations ago with an admixture proportion of 0.0225, where Neanderthals diverged 18,966 generations ago. The mutation and recombination rates were set to 1.25 × 10^−8^ and 1.0 × 10^−8^ per base pair per generation for ArchIE_3D19, 1.2 × 10^−8^ and 0.7 × 10^−8^ for BonoboGhost_4K19, and 1.29 × 10^−8^ and 1.0 × 10^−8^ for HumanNeanderthal_4G21, respectively. For each model, we simulated 50 diploid individuals from the reference population and 50 diploid individuals from the target population. To generate training data for logistic regression and extra-trees classifiers, we simulated sequences of 50,000 bp for ArchIE_3D19 and HumanNeanderthal_4G21, and 40,000 bp for BonoboGhost_4K19, assuming shorter introgressed fragments in this model. Each simulated sequence was labeled as introgressed if the introgressed proportion exceeded 0.7, labeled as non-introgressed if it was below 0.3, and discarded otherwise. To generate training data for UNet++, we simulated sequences of 100,000 bp, split them into genotype matrices with 192 variants, and upsampled the reference and target populations to 56 diploid individuals each. For testing, we simulated 200 Mb of sequence and processed the data using the same procedures as in training for each model. For each model, ten independent training runs were performed with different random seeds to assess robustness.

**Figure 1.**
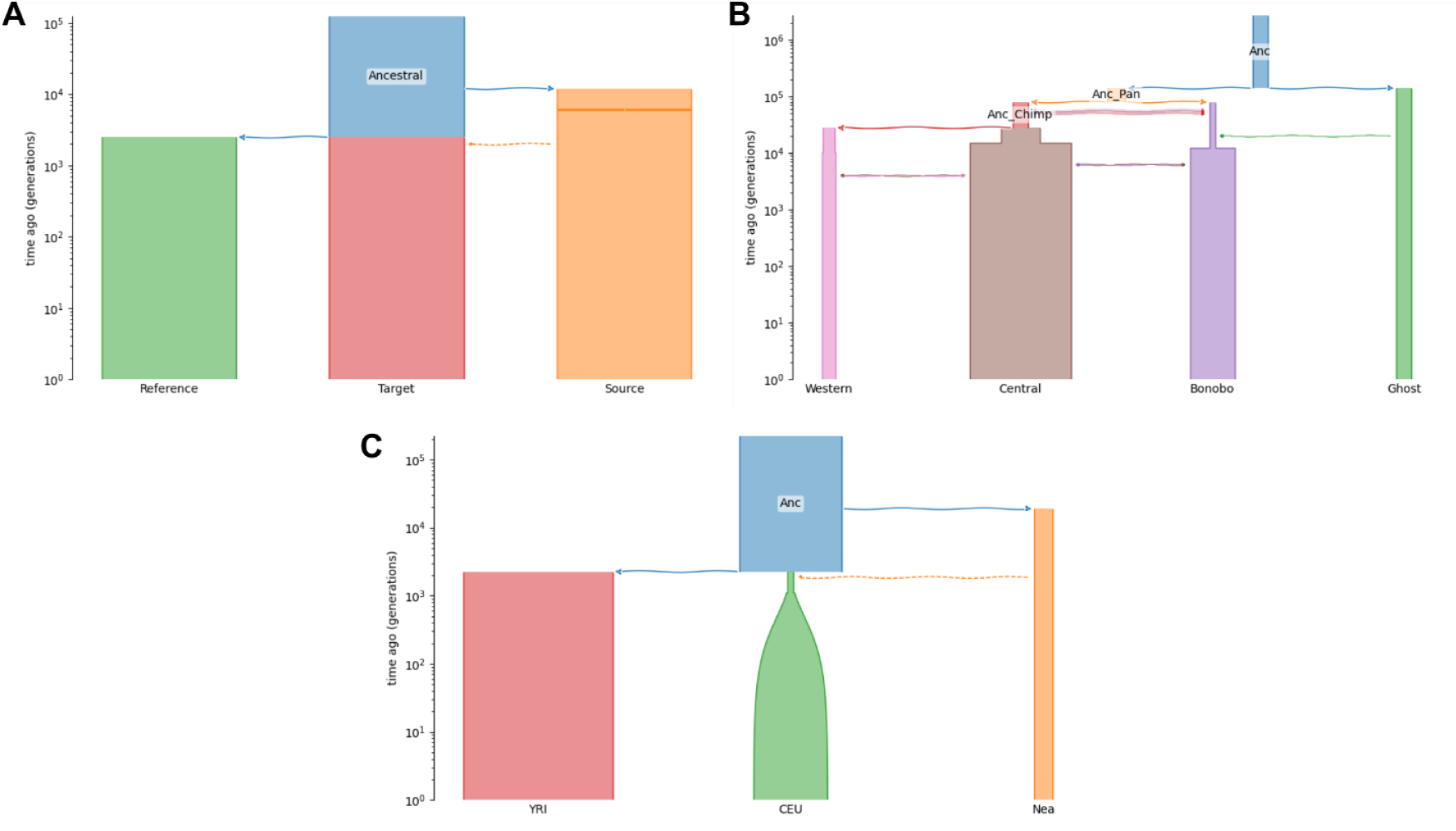
Demographic models used for performance evaluation. (A) ArchIE_3D19. (B) BonoboGhost_4K19. (C) HumanNeanderthal_4G21.

Because logistic regression and extra-trees classifiers produce segment-level introgression probabilities, whereas UNet++ produces allele-level probabilities, we evaluated all three machine learning models with precision-recall curves at both segment and allele levels, for both phased and unphased data (Figures 2 and 3). For segment-based evaluation, predictions from logistic regression and extra-trees classifiers were used directly as genomic segments. UNet++ predictions were converted to segment-level predictions by assigning variants to genomic windows with a length of 100,000 bp and averaging allele-level probabilities within each window and sample. Windows with probabilities above a given cutoff were called as introgressed, and adjacent positive windows from the same sample were merged into inferred tracts. For allele-based evaluation, UNet++ predictions were used directly. Predictions from logistic regression and extra-trees classifiers were converted to allele-level probabilities by projecting each sliding-window prediction onto the variant positions it covered and assigning each variant the mean probability across all overlapping windows.

**Figure 2.**
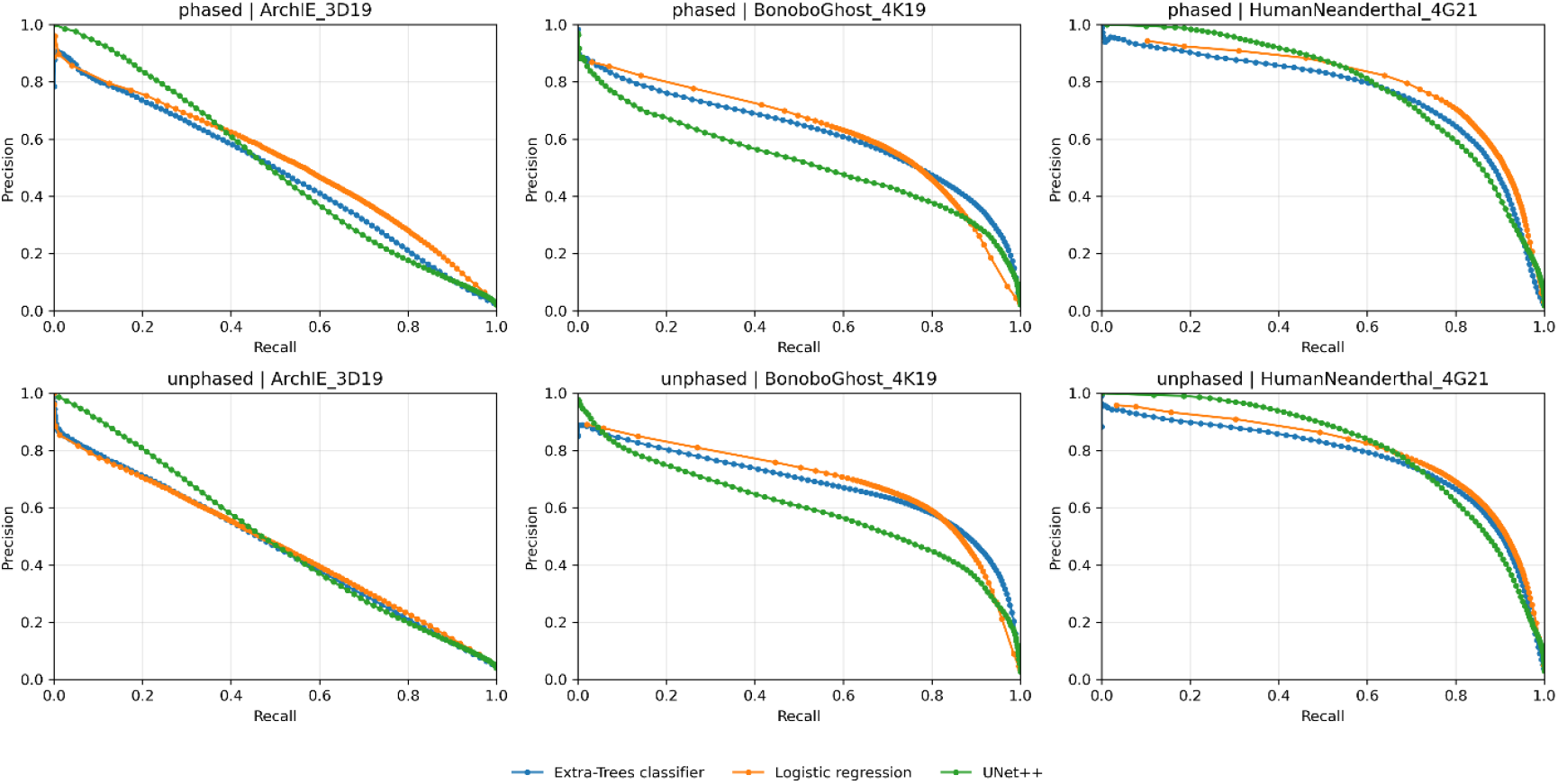
Precision–recall curves for machine learning models implemented in GAISHI using the segment-based evaluation. Each panel reflects performance for a given demographic model using phased or unphased data as input.

**Figure 3.**
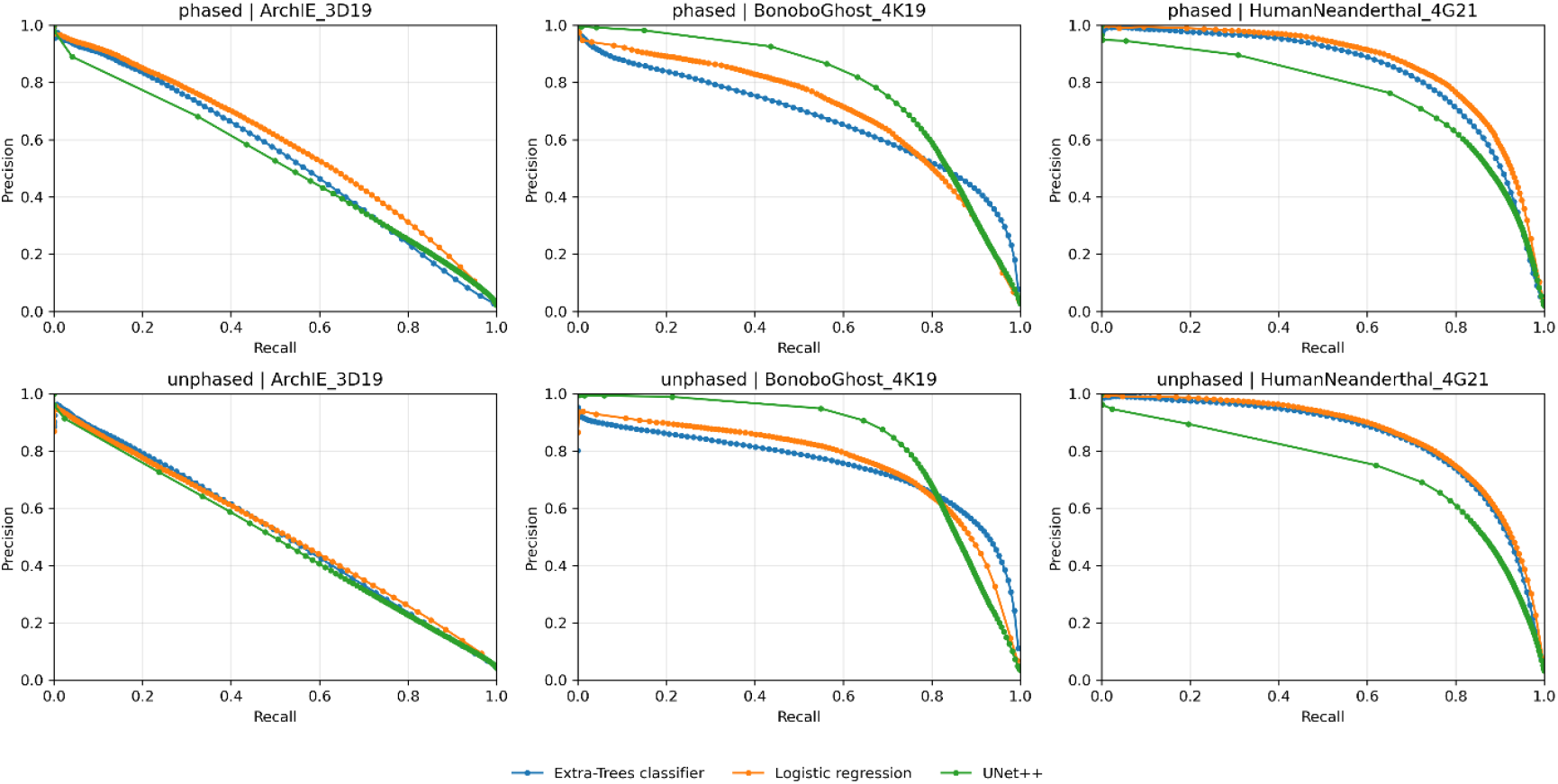
Precision–recall curves for machine learning models implemented in GAISHI using the allele-based evaluation. Each panel reflects performance for a given demographic model using phased or unphased data as input.

Our results suggest that, in the ArchIE_3D19 demographic model, all three machine learning models performed similarly, and their performance was the lowest among the three demographic models, indicating that this scenario is particularly challenging for confident inference of introgressed segments. In the BonoboGhost_4K19 demographic model, UNet++ achieved higher precision at recall > 0.8 than the other machine learning models in the allele-based evaluation. In the HumanNeanderthal_4G21 demographic model, all three machine learning models performed similarly in the segment-based evaluation, whereas logistic regression and extra-trees classifiers performed slightly better than UNet++ in the allele-based evaluation. This suggests that even though the UNet++-based method was designed to provide allele-based predictions, its performance compared to other algorithms depends on the demographic model. Overall, no single model performed best across all scenarios, indicating that users should select machine learning models according to their data and demographic model.

## Conclusion

GAISHI provides a flexible framework for detecting genomic footprints of ghost introgression using machine learning. Because the underlying machine learning models are general-purpose and can be adapted to different inference goals via task-specific features and labels, GAISHI has the potential to be extended to tasks such as adaptive introgression by incorporating other statistics, such as those implemented in the SAI package (Huang et al. 2025b). Moreover, given the rapid progress in machine learning, GAISHI can readily integrate additional model architectures as they emerge.

## Acknowledgements

The authors thank the Life Science Compute Cluster at the University of Vienna and the Multi-Site Computer Austria of the Austria Scientific Computing.

## Author contributions

X.H. designed the study. X.H. tested ArchIE. J.H. tested IntroUNET. X.H. implemented GAISHI. H.P. tested early versions of GAISHI. X.H. and M.K. wrote the manuscript.

